# Calpain-mediated proteolysis of vimentin filaments is augmented in Giant Axonal Neuropathy (GAN) fibroblasts exposed to hypotonic stress

**DOI:** 10.1101/2022.07.31.501244

**Authors:** Cassandra L. Phillips, Dong Fu, Laura E. Herring, Diane Armao, Natasha Snider

## Abstract

Giant Axonal Neuropathy (GAN) is a pediatric neurodegenerative disease caused by loss-of-function mutations in the E3 ubiquitin ligase adaptor gigaxonin, which is encoded by the *GAN* (*KLHL16*) gene. Gigaxonin regulates the degradation of multiple intermediate filament (IF) proteins, including neurofilaments, GFAP, and vimentin. In the absence of functional gigaxonin, GAN patients display abnormal cytoplasmic IF aggregates in multiple cell types. Understanding how normal IF networks and abnormal IF aggregates respond and are processed under physiologic stress can reveal new GAN disease mechanisms and potential targets for therapy. Here we tested the hypothesis that hypotonic stress-induced vimentin proteolysis is impaired in GAN. In both GAN and control fibroblasts exposed to hypotonic stress, we observed time-dependent vimentin cleavage, resulting in two prominent ~40-45 kDa fragments readily detectable by immunoblot in the total cell lysates and detergent-insoluble fractions. However, vimentin proteolysis occurred more rapidly and extensively in GAN cells compared to unaffected controls. Both fragments were generated earlier in GAN cells and at 4-6-fold higher levels (p<0.0001) compared to control fibroblasts. To test enzymatic involvement, we determined the expression levels and localization of the calcium-sensitive calpain proteases-1 and -2 and their endogenous inhibitor calpastatin. While the latter was not affected, the expression of both calpains was 2-fold higher in GAN cells compared to control cells (p<0.01). Moreover, pharmacologic inhibition of calpains with MDL-28170 or MG-132 attenuated vimentin cleavage, the latter resulting in >95% reduced cleavage (p<0.0001). Imaging analysis revealed striking colocalization between large perinuclear vimentin aggregates and calpain-2 in GAN fibroblasts. This colocalization was dramatically altered by hypotonic stress, where selective breakdown of IF networks with relative sparing of IF aggregates occurred rapidly in GAN cells and coincided with cytoplasmic redistribution of calpain-2. Finally, mass spectrometry-based proteomics revealed that phosphorylation at Ser-412, located at the junction between the central “rod” domain and C-terminal “tail” domain on vimentin, is involved in this stress response. Over-expression studies using phospho-deficient (S412A) and phospho-mimic (S412D) mutants revealed that Ser-412 is important for filament organization, solubility dynamics, and cleavage of vimentin upon hypotonic stress exposure. Collectively, our work reveals that osmotic stress induces calpain- and proteasome-mediated vimentin degradation and IF network breakdown. These effects are significantly augmented in the presence of disease-causing *KLHL16* mutations that alter IF spatial distribution and intermediate filament organization. While the specific roles of calpain-generated vimentin IF fragments in GAN cells remain to be defined, this proteolytic pathway is translationally-relevant to GAN because maintaining osmotic homeostasis is critical for nervous system function.

## Introduction

Intermediate filament (IF) proteins form filamentous networks that support cell structure and function [1]. Encoded by more than 70 individual genes in humans, IFs serve as organizers of the cytoplasmic space, scaffolds of interacting proteins within signaling networks, and mediators of stress responses [2]. With respect to the latter, IF protein networks are known to be extensively remodeled in cells undergoing stress, which is critical for their many cytoprotective functions [3; 4]. The remarkable plasticity of cytoplasmic IFs is highly dependent on various post-translational modifications (PTM), including phosphorylation, acetylation, sumoylation, and enzymatic proteolysis [5]. Regulated cross talk between the various PTMs on IF proteins impart a significant level of complexity to the system to ensure appropriate homeostatic and allostatic responses.

While IFs are critical for providing cells with stress resilience, chronic unresolved stress or genetic mutations can give rise to focal abnormal cytoplasmic IF accumulations (aggregates) in various cell types [6; 7; 8; 9]. The structural nature and precise roles of cell-type specific IF aggregates remain unclear. Yet, in the context of IF-associated human disease, gain of IF aggregates over time is accompanied by clinical decompensation and disease progression [10; 11; 12; 13; 14; 15]. This is particularly evident in the pediatric neurodegenerative disease Giant Axonal Neuropathy (GAN), which is caused by loss-of-function mutations in the gene *KLHL16* (also called *GAN)* [16]. *KLHL16* encodes the protein gigaxonin [17] - an adapter of an E3 ubiquitin-ligase complex that targets IF proteins for proteasomal degradation [18]. In the absence of functional gigaxonin, multiple IF proteins, including desmin, neurofilaments, GFAP, and vimentin, accumulate in different cell types in GAN patients [19; 20; 21; 22]. Pathologic diagnosis of GAN is based on dense bundles of IF accumulations causing focal, greatly enlarged, axonal swellings, or “giant axons,” after which the disease was named [23]. GAN affects both the peripheral nervous system (PNS) and the central nervous system (CNS). The natural history of GAN is characterized by progressive motor and sensory loss, with patients being non-ambulatory by the second decade of life, and death usually during the third decade [24]. Therefore, new mechanistic insights into the genesis and dismantling of IF aggregates will advance the development of therapeutic targets for IF-associated diseases like GAN. Importantly, proteolytic pathways for IF degradation in GAN patient cells exposed to external stress have not been characterized.

In general, exposure to stress affects the IF network composition and organization, as well as IF solubility dynamics, which are significantly altered in cells with persistent IF aggregates [25]. Osmotic stress in particular is known to regulate IF solubility and network formation [26]. Recently, it was shown that brief hypotonic exposure caused rapid and reversible reorganization and breakdown of the vimentin IFs, and that this occurred prior to any significant alterations in the actin and microtubule cytoskeletal networks [27]. Moreover, it is known that cells lacking vimentin are more susceptible to hypotonic stress, suggesting that rapid vimentin re-organization may be important in this cytoprotective response [26]. Osmotic stress plays an active role during inflammatory programmed cell death (pyroptosis) [28], where vimentin cleavage and loss of vimentin IF network leads to cell rupture and release of immunostimulatory cellular components. Given the importance of osmotic stress-dependent vimentin cleavage in different cellular responses, we tested the hypothesis that stress-induced vimentin proteolysis is impaired in GAN.

To that end, we exposed GAN patient-derived and unaffected control fibroblasts to hypotonic-stress for brief periods and analyzed vimentin changes biochemically and by immunofluorescence imaging. Collectively, our results show that hypotonic stress induces calpain- and proteasome-mediated vimentin degradation and IF network breakdown, and that there are regulatory phosphorylation sites located at the junction between the rod and C-terminal tail domain on vimentin that are involved in this IF stress response. Contrary to our hypothesis, vimentin breakdown occurred more rapidly and extensively in GAN cells compared to controls, raising the possibility that vimentin cleavage products may be involved in cellular dysfunction in GAN. While the specific roles of calpain-generated vimentin fragments in GAN remain to be defined, this proteolytic pathway is translationally-relevant to the disease pathogenesis of GAN because maintaining osmotic homeostasis is critically important for homeostasis and nervous system function [29].

## Results

### Hypotonic stress promotes time-dependent cleavage and reorganization of vimentin IFs in GAN fibroblasts

Given the different organization of vimentin IFs in GAN fibroblasts, which is characterized by the presence of both cytoplasmic filaments and perinuclear ovoid bundles [30], we asked whether vimentin cleavage will occur in a similar fashion in GAN cells. To that end, we exposed GAN patient-derived fibroblasts to hypotonic stress using water exposure, as was done previously [27] for 0-8 minutes. Since osmotic stress is known to alter IF protein solubility, we compared vimentin in total lysates, Triton X detergent-insoluble, and detergent-soluble fractions. As shown in **Fig. 1A**, there was time-dependent vimentin cleavage in the GAN fibroblasts, resulting in two prominent fragments (FR1, FR2) detected by western blot in the total cell lysates and detergent-insoluble fractions, in addition to full length vimentin. Small amounts of cleaved vimentin (FR1) were also seen in the detergent-soluble fraction after 8 minutes of treatment **(Fig. 1A)**. The two cleaved products were most abundant in the 8-minute exposure condition, indicating that prolonged exposure resulted in increased breakdown of vimentin and/or decreased downstream processing of the smaller fragments (**Fig. 1A**). Disruption of vimentin IFs in GAN fibroblasts was also observed by immunofluorescence imaging, as longer exposure time to hypotonic stress resulted in the progressive reduction of cytoplasmic vimentin filaments, while perinuclear aggregates persisted across conditions (**Fig. 1B**). These results indicated that vimentin IFs in GAN cells are sensitive to hypotonic stress, but the vimentin aggregates are more resistant to this treatment.

**Figure 1.**
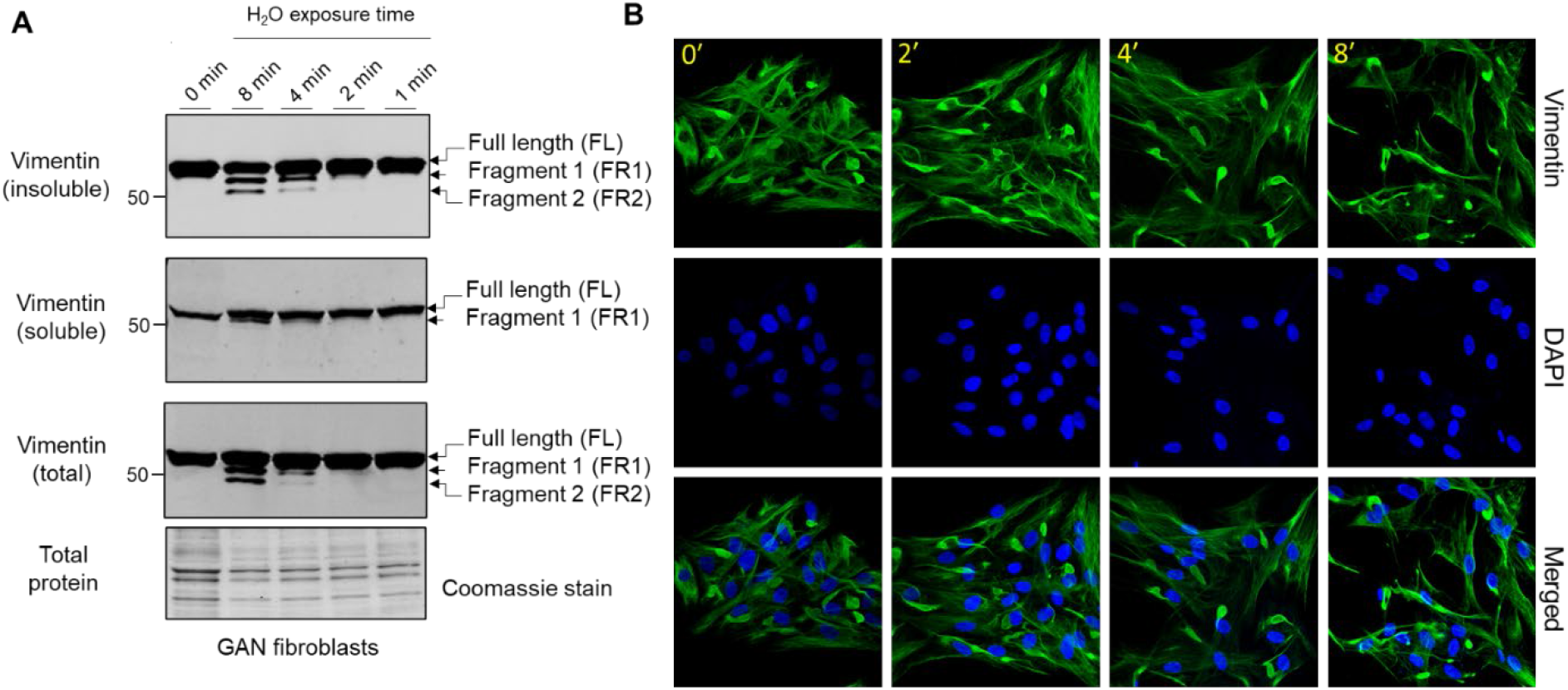
Hypotonic stress induces time-dependent cleavage and reorganization of vimentin IFs in GAN fibroblasts. **(A)** Immunoblotting of vimentin in reduced Triton-X insoluble pellet fractions (top), Triton-X soluble fractions (middle), and total cell (bottom) lysates from GAN fibroblasts exposed to hypotonic stress conditions (sterile molecular grade H_2_O) for 0-8 minutes. Only full length (FL) vimentin is present in the 0-minute condition (lane 1), but a time-dependent, osmotic stress-induced increase in vimentin cleavage is observed starting in the 2-minute condition (lane 4). The two vimentin fragments (FR1, FR2) produced are most abundant in the 8-minute condition (lane 2). Coomassie stain was used as a control for total protein. **(B)** Immunofluorescence analysis of GAN fibroblasts stained with vimentin (green) and DAPI (blue) after exposure to water for 0-8 minutes. Note the persistence of vimentin aggregates, but a reduction in vimentin filaments.

### Hypotonic stress-induced vimentin cleavage is augmented in GAN compared to control fibroblasts

Next, we asked whether vimentin cleavage under hypotonic stress is altered in GAN compared to control (unaffected) fibroblasts. Direct comparison of cleaved vimentin products over time revealed the presence of vimentin fragments at earlier time points and at higher levels in the detergent-insoluble fractions of GAN cells compared to control cells (**Fig. 2A**). In the GAN cells, both cleaved vimentin fragments can be observed in the 2-minute condition and were highly abundant in the 8-minute condition (**Fig. 2A**). In the control cells, there was no evidence of vimentin cleavage until the 8-minute condition, and the levels of the fragments were lower when normalized to full length vimentin **(Fig. 2B)**. Immunofluorescence staining for vimentin corroborated the biochemical analysis, indicating that vimentin IFs in control fibroblasts were significantly more resistant to re-organization and filament breakdown under hypotonic stress (**Fig. 2C**). These data suggested that vimentin IFs in GAN fibroblasts were more sensitive to hypotonic stress-induced cleavage. This prompted us to determine if calpains, proteolytic enzymes that have been previously implicated in vimentin cleavage, are involved in this vimentin cleavage pathway and if their expression is altered in GAN cells.

**Figure 2.**
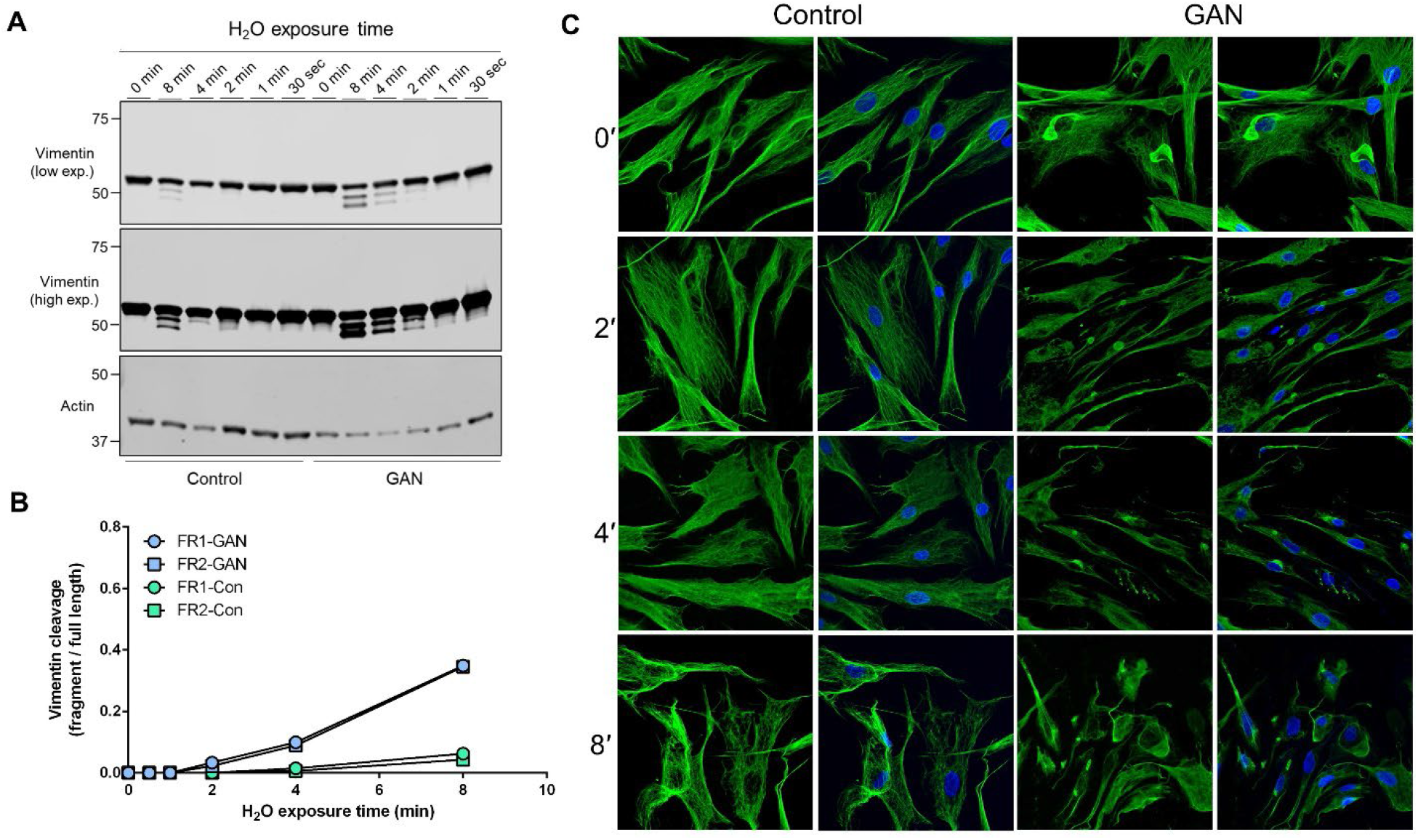
Hypotonic stress-induced vimentin cleavage is augmented in GAN compared to control fibroblasts. **(A)** Immunoblotting of vimentin (top and middle) and actin (bottom; loading control) in reduced Triton-X insoluble pellet fractions from control (lanes 1-6) and GAN (lanes 7-12) fibroblasts exposed to hypotonic stress conditions (sterile molecular grade H_2_O) for 0-8 minutes. Expression of the two vimentin fragments (FR1 and FR2) is increased in GAN patient cells compared to control cells. Low (top) and high (middle) exposure blots of the same membrane are shown for vimentin. **(B)** Densitometry quantification of vimentin fragment (FR) intensities relative to full length vimentin in control and GAN fibroblasts exposed to hypotonic stress for 0-8 minutes. Data points represent values from a single experiment and are representative of at least 3 independent experiments. **(C)** Immunofluorescence analysis of control and GAN fibroblasts stained with vimentin (green) and DAPI (blue) after exposure to water for 0-8 minutes. Note significant breakdown of the vimentin filament network in the GAN cells at an earlier timepoint (2’) compared to the control cells (8’).

### Vimentin cleavage is blocked by inhibitors of the proteasome and calpains, which are elevated in GAN

Pre-treatment with the calpain inhibitor MDL-28170, followed by 8 minutes of water exposure reduced, but did not eliminate vimentin cleavage in GAN cells (**Fig. 3A**). The fragments were nearly undetectable in the presence of MG-132, which inhibits calpains and the proteasome **(Fig. 3A).** Similarly, MG-132 eliminated vimentin cleavage in control fibroblasts **(Fig. 3B)**, suggesting involvement of similar proteolytic pathways. However, we detected significantly higher levels of calpain-1 and −2 in GAN cells compared to control cells, but no major differences in calpastatin, an endogenous inhibitor of calpain activity in cells **(Fig. 3B)**. The increased calpain expression correlated with increased cleavage of vimentin in GAN cells **(Fig. 3C)**. Therefore, increased calpain-1/2 expression may account, at least in part, for the increased vimentin cleavage in GAN cells exposed to hypotonic stress.

**Figure 3.**
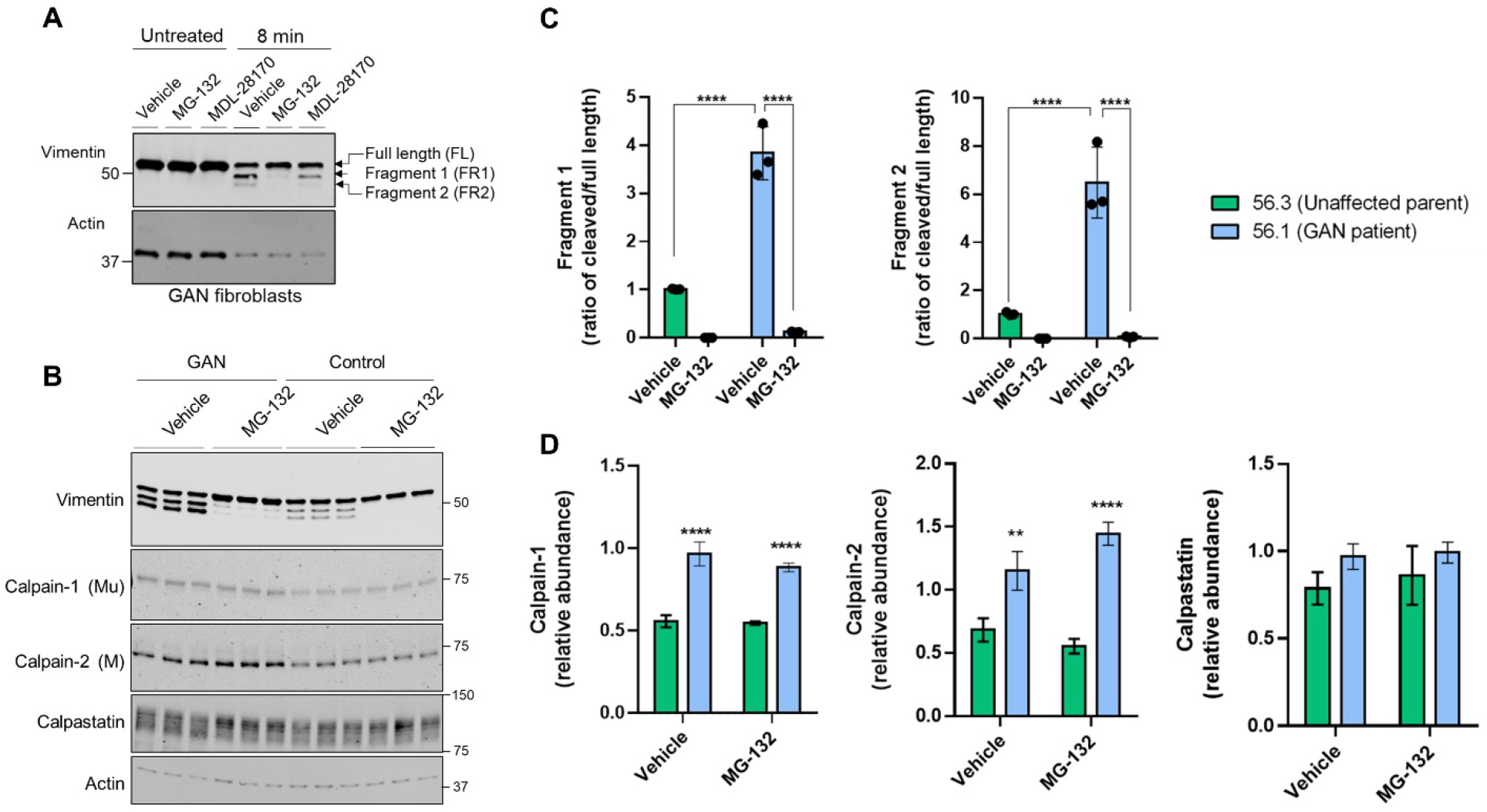
Vimentin cleavage is blocked by inhibitors of the proteasome and calpains, which are elevated in GAN. **(A)** Immunoblotting for vimentin (top) and actin (bottom; loading control) in Triton-X insoluble pellet fractions from untreated or water-exposed GAN fibroblasts in the absence or presence of the DMSO vehicle control, calpain inhibitor (MDL-28170), or calpain/proteasome inhibitor (MG-132). Note partial cleavage inhibition by MDL and near-complete inhibition by MG-132. **(B)** Immunoblotting for vimentin (top), calpain-1 (middle top), calpain-2 (middle), calpastatin (middle bottom), and actin (bottom; loading control) in reduced Triton-X insoluble and soluble fractions from GAN and control fibroblasts exposed to 8 minutes of hypotonic stress and treated with the DMSO vehicle control or MG-132. **(C)** Quantification of the relative fragment 1 (FR1) and fragment 2 (FR2) band intensities normalized to the full-length vimentin band for each cell line and treatment group (from panel B). ****p<0.0001; two-way ANOVA. **(D)** Quantification of calpain-1, calpain-2, and calpastatin levels from panel B. **p<0.01; ****p<0.0001; two-way ANOVA.

### Calpain-2 colocalizes with perinuclear vimentin aggregates and undergoes cytoplasmic redistribution upon hypotonic stress exposure

In addition to the biochemical analyses, we investigated calpain localization via immunofluorescence imaging. Interestingly, the imaging analyses revealed prominent colocalization between large perinuclear vimentin aggregates and calpain-2 in the untreated GAN patient fibroblasts (**Fig. 4A**). However, calpain-2 localization was dramatically altered following 8 minutes of hypotonic stress exposure (**Fig. 4B**). In addition to vimentin network breakdown in response to stress, calpain-2 staining became more punctate and distributed throughout the cytoplasm, which was largely devoid of filaments (**Fig. 4B**). This cytoplasmic redistribution of calpain-2 indicated a rapid and selective breakdown of filaments over aggregates in the GAN cells, which provided further evidence for the idea that calpain proteins are involved in this rapid, stress-induced degradation mechanism. The mechanisms by which calpain-2 associates with the perinuclear vimentin aggregates in GAN cells remain to be defined. We hypothesized that hypotonic stress-induced phosphorylation of vimentin may play a role in the release of calpain-2 from the aggregates, thereby leading to increased vimentin cleavage within the cytoplasmic network.

**Figure 4.**
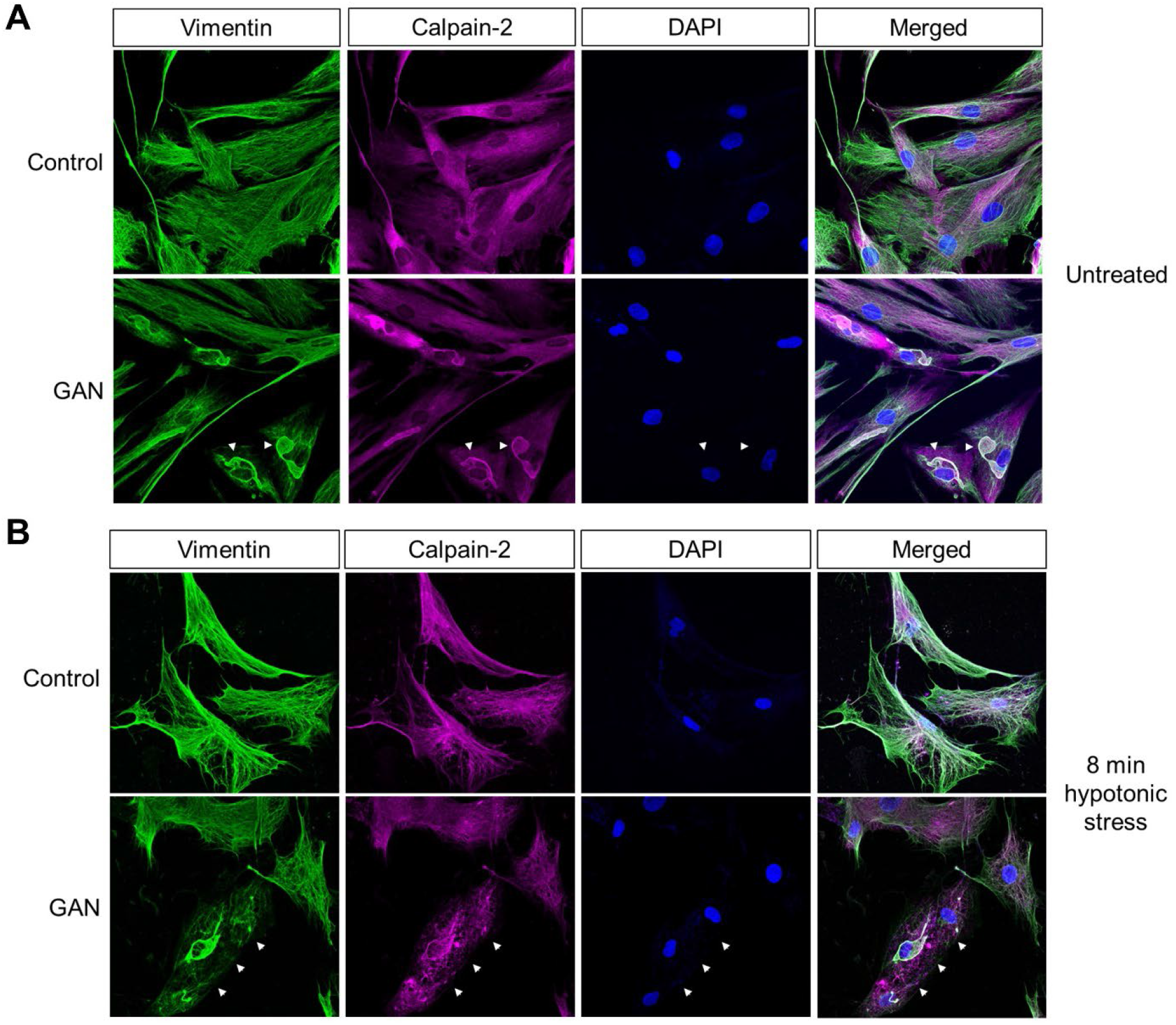
Calpain-2 colocalizes with perinuclear vimentin aggregates and undergoes cytoplasmic redistribution upon hypotonic stress exposure. **(A)** Calpain colocalizes with vimentin aggregates in untreated GAN fibroblasts. Immunofluorescence analysis of untreated control (top) and GAN (bottom) fibroblasts stained with vimentin (green), calpain-2 (magenta), and DAPI (blue). Large perinuclear vimentin aggregates (green) colocalize with calpain-2 (magenta) in GAN patient cells (white arrowheads). **(B)** Calpain redistribution and vimentin network breakdown under hypotonic stress. Immunofluorescence analysis of control (top) and GAN (bottom) fibroblasts exposed to hypotonic stress for 8 minutes and stained with vimentin (green), calpain-2 (magenta), and DAPI (blue). Calpain-2 (magenta) becomes more punctate and localized to vimentin filaments, indicating the selective loss of the vimentin filament network as compared to the relative preservation of vimentin aggregates (green) in GAN fibroblasts (white arrowheads).

### Site-specific and time-dependent dephosphorylation in response to hypotonic stress regulates vimentin solubility

Phosphorylation sites on vimentin isolated from untreated or hypotonic stress-exposed GAN fibroblasts were mapped by mass spectrometry analysis **(Fig. 5A)**. Somewhat surprisingly, we did not observe a significant increase in vimentin phosphorylation at the sites identified in the head (S29, S39, S42, S56), rod (Y276), and tail (S409/412, T426, S430) domains. However, we noted a highly time-dependent decrease in S409/412 phosphorylation that was statistically significant and unique among the sites that were identified in the analysis (**Fig. 5A**). Moreover, predictive analysis of calpain cleavage sites on vimentin using DeepCalpain identified cleavage within peptide 411-426, which contains the pSer412 site and the 418-419 predicted calpain-targeted junction **(Fig. 5B)**. Based on AlphaFold analysis, this serine residue marks the end of the coiled-coil rod domain **(Fig. 5C)**, suggesting that it likely has an important role in filament dynamics. To test that, we made phospho-mimic **(S409D, S412D)** and phospho-deficient **(S409A, S412A)** mutants and determined how they affected vimentin cleavage. Using BHK-21 fibroblasts, we over-expressed WT and mutant vimentin for 24 hours and induced cleavage by water treatment. Consistent with the proteomic analysis, we observed increased cleavage in the phospho-deficient S412A mutant **(Fig. 5D)**, albeit not statistically significant **(Fig. 5E)**. However, in the case of over-expressed vimentin, the fragments were detected primarily in the soluble fraction **(Fig. 5D)** and solubility was significantly decreased in the S412D phospho-mimic mutant **(Fig. 5F)**. In agreement with the reduced solubility, the S412D mutant exhibited abnormal filament structure with evidence of bundling, whereas S412A displayed bundles and normal filaments (**Fig. 5G**). Of note, there were no changes in filament solubility or organization when Ser-409 was similarly mutated (not shown). Combined, these data demonstrated that Ser-412 is an important site for vimentin filament assembly and filament regulation during hypotonic stress exposure.

**Figure 5.**
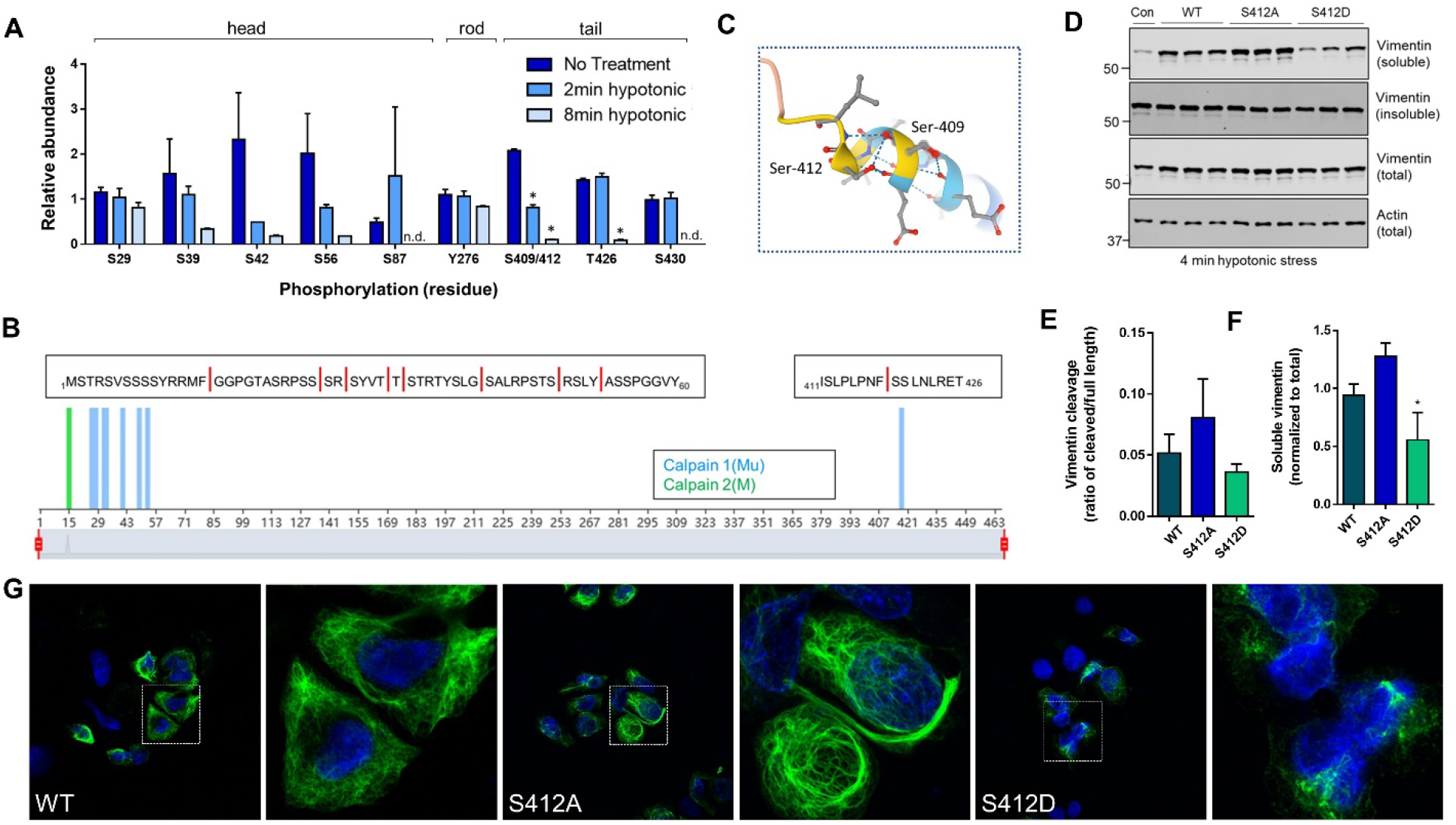
Site-specific and time-dependent dephosphorylation in response to hypotonic stress regulates vimentin solubility. **(A)** Mass spectrometry analysis for phosphorylation site changes on vimentin in GAN fibroblasts after 2 minutes and 8 minutes of hypotonic stress exposure. *p<0.001. Multiple t-tests. **(B)** Predictive analysis of calpain cleavage sites on vimentin using the DeepCalpain tool. Peptide sequences span the N-terminal head domain (aa1-60), containing 8 predicted cut sites (red lines) and C-terminal tail domain peptide (aa411-426) containing a single predicted cut site. **(C)** Model of vimentin rod/tail junction (AlphaFold) with the localization of Ser-409/412. **(D)** Immunoblotting for vimentin (top, middle) and actin (bottom; loading control) in Triton-X soluble (top), Triton-X insoluble (middle top), and total cell (middle bottom, bottom) lysates from BHK-21 cells transfected with WT, S412A, and S412D constructs and exposed to hypotonic stress for 4 minutes. **(E-F)** Quantification of vimentin solubility (E) and cleavage (F) in WT and mutant vimentin. *p<0.05; one-way ANOVA. **(G)** Immunofluorescence analysis of SW13 Vim^-^ cells transfected with WT vimentin (left) and vimentin mutants S412 A (middle) and S412D (right) and stained with vimentin (green) following a 24-hour transfection period. White squares indicate magnified areas.

## Discussion

In this study, we demonstrated that hypotonic stress exposure promoted rapid vimentin proteolysis and IF network breakdown in GAN patient-derived and unaffected control fibroblasts. Surprisingly, we found that vimentin IFs in GAN fibroblasts were more sensitive to hypotonic stress-induced cleavage, suggesting that the presence of *KLHL16* mutations may be associated with increased IF fragmentation under stress. Based on pharmacological inhibition experiments, we concluded that this proteolytic pathway was a calpain- and proteasome-mediated process. Whether proteolytic vimentin fragments have distinct functions and whether they regulate acute and chronic cellular stress response and recovery remains to be elucidated. We speculate that any potential biological function of the fragments will likely be determined by their stability in the cell. Many calpain-generated fragments are short-lived because they are substrates for the N-end rule pathway, a ubiquitin-proteasome dependent system that recognizes destabilizing N-terminal residues (Arg, Lys, His, Leu, Trp, Phe, Tyr, and Ile), which are directly recognized by N-recognins [31]. However, most of the predicted calpain cut sites on vimentin expose an N-terminal residue on the resulting fragment that would be highly stabilizing (half-life >30hr) [32]; including the putative head (N-terminal Ser, Ala, Gly), and tail (N-terminal Ser) domain fragments (**Fig. 5B**). Based on such observations, we hypothesize that the vimentin fragments will be long-lived and may exert significant impact on cellular function. This will be tested in future work, particularly in the context of GAN disease-relevant cell types harboring abnormal IF aggregates, specifically neurons and astrocytes.

Our imaging analysis showed striking colocalization between perinuclear vimentin aggregates and calpain-2, which had significantly increased expression in the GAN patient fibroblasts. While the reason for the increased calpain expression in GAN fibroblasts remains to be investigated, it is possible that association with vimentin aggregates promotes calpain stability. Active calpain-2 has been localized to protein aggregates associated with other neurodegenerative diseases, such as hyper-phosphorylated Tau protein [33]. In that regard, GAN may share similar mechanisms to other neurodegenerative disorders. In Alzheimer’s disease (AD), calpain proteases may be involved in the proteolysis of amyloid precursor protein (APP), which is involved in the formation of amyloid plaques [34]. Interestingly, AD mice treated with calpastatin to inhibit calpain activity exhibited improved cognitive function and synaptic transmission [35]. Additionally, treatment with calpastatin or other calpain inhibitors mitigated dopaminergic neuron loss in mice with Parkinson’s disease (PD) [36]. Furthermore, calpain-mediated cleavage of TDP-43 in motor neurons contributes to the aggregates observed in Amyotrophic lateral sclerosis (ALS) [37].

Increased levels of activated calpains have been detected in postmortem tissue from the prefrontal cortex and caudate nucleus of patients with AD and Huntington’s disease (HD) respectively [38; 39]. Since calpain-mediated cleavage is implicated in multiple neurodegenerative diseases that are characterized by abnormal protein aggregation, it is plausible that calpain proteases may also contribute to the pathologic IF accumulations observed in GAN through cleavage of IF proteins. This mechanism warrants investigation in future studies to determine if there are overlapping cellular mechanisms between GAN and other neurodegenerative diseases known to exhibit elevated calpain activity [34].

Colocalization between vimentin and calpain-2 was significantly altered by hypotonic stress exposure, where selective breakdown of cytoplasmic IF networks with relative sparing of IF aggregates occurred rapidly in the patient cells and coincided with cytoplasmic redistribution of calpain-2. If, similar to the reported calpain-2/Tau interactions, calpain-2 associates with hyper-phosphorylated vimentin within the perinuclear aggregate, then decreased phosphorylation of vimentin upon stress induction may sever that interaction. Post-translational modifications (PTMs), particularly phosphorylation, are known to regulate IF functions and properties, including mediating IF stress responses [5]. PTMs can serve as a precursor to additional responses, such as reorganization or degradation during or following a stress event. Our mass spectrometry proteomics indicated that phosphorylation at Ser-412 on vimentin is involved in this hypotonic stress response. This residue has not been well-characterized in the literature, although some groups have observed phosphorylation at residue S412 under physiological conditions [5; 40]. Recently, it was shown that modifications to S412 may influence filament assembly or positioning of the tail domain [40]. Under normal conditions, an antibody targeting the 411-423 epitope of vimentin readily detected filaments throughout the network, whereas another epitope localized to the 419-438 segment was less accessible except for areas with rarefied filaments due to a compacted formation of the tail [40]. Interestingly, both epitopes were similarly accessible with a phospho-mimic S412 vimentin mutant, suggesting that the tail domain was in a more open formation as a result of the mutation [40]. Those findings, along with our data, support the idea that Ser-412 regulates vimentin filament organization, solubility, and degradation. Future efforts will be dedicated to further probing the relationship between key regulatory PTMs sites and proteolytic fragmentation of vimentin in response to osmotic stress.

Astrocytes are key players in the maintenance of cell volume homeostasis in the brain since they oversee fluxes of ions, water and osmolytes at their homeostatic concentrations [41]. Astrocyte IFs, including vimentin, are critical structures for cellular stress responses because astrocytes are sensitive to stress exposure, including osmotic and mechanical stresses resulting from ischemia, trauma, and brain edema [42; 43; 44]. The responsive nature of these cells can result in a *“reactive”* astrocyte phenotype, which may be beneficial during acute stress, but detrimental with chronic and prolonged stress [45]. Reactive astrocytes are prominent in GAN and other neurodegenerative disorders. Additionally, GAN astrocytes exhibit striking GFAP aggregates that co-accumulate with vimentin. Currently, it is unclear how abnormal cytoplasmic IF aggregates affect GAN astrocyte function, nor is it clear how impaired astrocyte function contributes to the disease pathology observed in GAN. Further, it remains unknown how GAN astrocyte IF aggregates and IF cytoplasmic networks respond under conditions of cellular stress. Our work described herein can serve as a springboard to address these questions in future studies.

## Methods

### Antibodies

The following primary antibodies and concentrations were utilized: rabbit anti-Vimentin (Cell Signaling Technology, D21H3, WB 1:1000, IF 1:200), mouse anti-Vimentin (Invitrogen, V9, WB 1:1000-1:2500, IF 1:100), mouse anti-Actin (NeoMarkers, ACTN05, WB 1:1000), mouse anti-Actin (Santa Cruz, SPM161, WB 1:1000), rabbit anti-Calpastatin (Cell Signaling Technology, WB 1:1000), rabbit anti-Calpain 1 (Cell Signaling Technology, large subunit Mu-type, WB 1:1000), rabbit anti-Calpain 2 (Cell Signaling Technology, large subunit M-type, WB 1:1000), and rabbit anti-Calpain 2 (Cell Signaling Technology, large subunit M-type, E3M6E, IF 1:200). The following secondary antibodies and concentrations were utilized: IRDye 800CW goat anti-rabbit IgG (LI-COR, WB 1:5000), IRDye 680RD donkey anti-mouse IgG (LI-COR, WB 1:5000), and Alexa 488- and Alexa 568-conjugated goat anti-mouse and anti-rabbit antibodies (Invitrogen, IF 1:500).

### Cell lines

GAN patient-derived and normal control skin fibroblasts were provided by Dr. Steven Gray. The cells were thawed initially in DMEM (Gibco) in 20% FBS (GenClone, Lot: P093156) and 1% penicillin/streptomycin (Thermo Fisher Scientific) and passaged every 3-4 days with 0.05% Trypsin-EDTA (Gibco), and after a few passages, the cells were maintained in 10% FBS media. SW13 vimentin negative (Vim^-^) cells and BHK-21 cells were thawed and maintained in DMEM (Gibco) in 10% FBS (GenClone, Lot: P093156) and 1% penicillin/streptomycin (Thermo Fisher Scientific) and passaged every 3-4 days with 0.25% Trypsin-EDTA (Gibco).

### Hypot onic stress experiments

For hypotonic stress experiments, the cells were plated on 6-well plates and treated with sterile molecular grade water (Corning). Following aspiration of media, the cells were treated 3mL/well of water and incubated in a 37°C incubator for the designated timepoint (i.e., 0 min, 0.5 min, 1 min, 2 min, 4 min, or 8 min). After treatment, the water was aspirated and Triton X-100 buffer (Triton X-100, 0.5 M EDTA, PBS, ddH_2_O) with phosphatase (Roche) and protease (Roche) inhibitors was added directly to each well. With the plate on ice, the cells were scraped from the wells and collected into Eppendorf tubes. At this stage, total cell lysate (TCL) samples were prepared by transferring some of the sample to a new tube and adding an equal volume of 2X Novex™ Tris-Glycine SDS Sample Buffer (Thermo Fisher Scientific). The remaining samples were spun down in a tabletop centrifuge at 14,000 RCF at 4°C to separate the detergent-soluble and insoluble fractions. The supernatant of the samples was transferred to a new tube, and for the soluble fraction, a smaller volume was then aliquoted to a different tube with an equal volume of SDS sample buffer. The remaining pellet was resuspended in SDS sample buffer to make the insoluble fraction. All samples with SDS sample buffer were heated at 95°C for 5 minutes and reduced with 5% 2-mercaptoethanol (Sigma) as needed.

### Pharmacological inhibition of calpain proteins

*MDL-28170 (Cell Signaling Technology):* A 100mM MDL-28170 stock solution was made by dissolving the entire vial (5mg) in DMSO. GAN patient and control fibroblasts were plated on 6-well plates, then pre-treated for 1 hour in a 37°C incubator with the inhibitor or the DMSO vehicle control; the final inhibitor concentration of 100uM/well (based on IC_50_ value of 10uM) was achieved by adding 500uL of inhibitor solution to 2.5mL media already in the wells. Following pre-treatment, cells were treated with either the inhibitor or the DMSO vehicle control both with and without sterile molecular grade water; the final inhibitor concentration of 100 uM/well was achieved by adding 3mL/well of the inhibitor/water (hypotonic treatment) or inhibitor/media (no treatment) solutions. The cells were exposed to hypotonic stress for 8 minutes (+/- inhibitor) in a 37°C incubator, then harvested and separated into TCL, detergent-soluble, and detergent-insoluble fractions as described above. *MG-132 (Cell Signaling Technology):* A 10mM MG-132 stock solution was made by dissolving the entire vial (1mg) in DMSO. GAN patient and control fibroblasts were plated on 6-well plates, then pre-treated for 1 hour in a 37°C incubator with the inhibitor or the DMSO vehicle control; the final inhibitor concentration of 10uM/well (based on IC_50_ value of 1.25uM) was achieved by adding 500uL of inhibitor solution to 2.5mL media already in the wells. Following pre-treatment, cells were treated with either the inhibitor or the DMSO vehicle control both with and without sterile molecular grade water; the final inhibitor concentration of 10 uM/well was achieved by adding 3mL/well of the inhibitor/water (hypotonic treatment) or inhibitor/media (no treatment) solutions. The cells were exposed to hypotonic stress for 8 minutes (+/- inhibitor) in a 37°C incubator, then harvested and separated into TCL, detergent soluble, and detergent insoluble fractions as described above.

### Site-directed mutagenesis and transfection of vimentin mutants

Mutagenesis of vimentin (pCMV6-XL5 vector; Origene) was conducted with the QuikChange II Site-Directed Mutagenesis Kit (Aligent Technologies) to generate the following point mutations: S409A, S409D, S409E, S412A, S412D, and S412E. Sanger sequencing for the entire vimentin coding sequence was completed to verify the point mutations were present in the sequence without off-target changes. For transfection experiments for immunofluorescence imaging, SW13 Vim^-^ cells were seeded onto 4-well chamber slides and transfected with vimentin wild-type and mutant plasmids along with lipofectamine 2000 that was utilized in accordance with product instructions (Thermo Fisher Scientific). Media was changed for the cells 6 hours after transfection, and cells were fixed at the 24-hour timepoint (see below for details on immunofluorescence staining procedures). For transfection and hypotonic stress experiments, BHK-21 cells were plated on 6-well plates and transfected with vimentin wild-type and mutant plasmids plus lipofectamine 2000. Media was changed for the cells 6 hours after transfection, and cells were treated with sterile molecular grade water at the 24-hour timepoint. The cells were treated with hypotonic stress for 4 minutes in a 37°C incubator, then harvested and separated into TCL, detergent soluble, and detergent insoluble fractions as described above.

### Preparation of protein lysates and immunoblotting

For immunoblotting, samples were separated on 10% or 4-20% gradient Novex™ WedgeWell™ Tris-Glycine gels (Thermo Fisher Scientific) for 40 minutes at 225V and transferred at 40V overnight at 4°C onto nitrocellulose membranes. Gels were stained with Coomassie following each transfer to verify normalization. The membranes were blocked in 5% non-fat milk (NFM) dissolved into 0.1% tween 20/PBS (PBST) at room temperature for 30 minutes. The membranes were incubated in primary antibodies diluted in 5% NFM/PBST at room temperature for 1 hour or at 4°C overnight (see concentrations above), then washed 3x with PBST for 5 minutes each. The membranes were incubated with secondary antibodies diluted in 5% NFM/PBST at room temperature for 1 hour (see concentrations above), washed 3x with PBST and 1x with PBS for 5 minutes each, then scanned with a LI-COR Odyssey CLx machine. Protein lysates were normalized by either method: staining gel with Coomassie before running the western blots (densitometry conducted with Adobe Photoshop), or blotting for pan-actin and normalizing the bands of interest to the pan-actin band intensity (densitometry conducted with Image Studio version 5.2).

### Immunofluorescence, imaging, and analysis

Cells were fixed in methanol at −20°C for 15 minutes, washed 2-3x with PBS for 5 minutes each, and blocked in Buffer B (2.5% Bovine Serum Albumin (Sigma), 2% normal goat serum (Gibco), PBS) at room temperature for 1 hour. Cells were incubated with primary antibodies (see concentrations above) at room temperature for 2 hours, followed by 3x 5-minute PBS washes, then incubated with Alexa Fluor-conjugated secondary antibodies (see concentrations above) at room temperature for 1 hour and washed 3x with PBS for 5 minutes each. Finally, cells were incubated in DAPI (Invitrogen), washed 3x with PBS for 5 minutes each, and mounted in Fluoromount-G (SouthernBiotech) overnight. Cells were imaged on Zeiss 880 confocal laser scanning microscope using a Plan-Neofluar 40x/1.3 oil WD0.21objective.

### Mass spectrometry

*Sample preparation:* GAN patient fibroblasts were plated on 10cm plates, then treated with 10mL/plate sterile molecular grade water in a 37°C incubator at designated timepoints (0 min, 2 min, and 8 min). After treatment, the water was aspirated and Triton X-100 buffer (see above for details) was added directly to each plate. With the plate on ice, the cells were scraped from the wells and collected into Eppendorf tubes, spun in a tabletop centrifuge at 14,000 RCF at 4°C, then the supernatant of the samples was transferred to a new tube. High salt buffer (KCL, NaCl, Tris-HCl, 0.5M EDTA, Triton X-100, ddH_2_O) with phosphatase (Roche) and protease (Roche) inhibitors was added to the remaining pellets, the samples were thoroughly dounced, spun in a tabletop centrifuge at 14,000 RCF at 4°C, then the sample supernatant was transferred to a new tube. 1x PBS/EDTA 5mM was added to the remaining pellets, which were vortexed and spun at 14,000 RCF at 4°C. The sample supernatant was transferred to a new tube and the remaining pellets were resuspended in SDS sample buffer to make insoluble fractions, which were then heated at 95°C for 5 minutes. Insoluble fraction samples were loaded in a 4-20% gradient gel and run for about 40 minutes at 225V, then the gel was stained with GelCode™ Blue Stain for 1 hour at room temperature and de-stained overnight in DDI H_2_O. The gel bands of interest were extracted for mass spectrometry analysis, and the excised bands were reduced, alkylated, and digested overnight at 37°C with trypsin. The extracted peptides were desalted using C18 ZipTips. *LC/MS/MS analysis and data analysis*: The samples were analyzed in duplicate on an Easy-nLC™ 1200-QExactive HF system (Thermo Fisher Scientific) as previously described [46], and the raw data was analyzed in Proteome Discoverer v2.5 (Thermo Fisher Scientific). The data were searched against the Uniprot human database appended with a database of common contaminants, and tryptic peptides were identified using the following parameters: precursor mass tolerance was set to 10 ppm, fragment mass tolerance was set to 20 ppm, and up to two missed cleavage sites were allowed. The variable modifications were set to phosphorylation of Ser, Thr, and Tyr, and oxidation of methionine. The ptmRS node was used to localize phosphorylation sites. The false discovery rate (FDR) was set to 1% and used to filter all data.

### Statistics

Image Studio version 5.2 (LI-COR) was used to perform densitometry on immunoblots, and Adobe Photoshop was used for densitometry on gels stained with Coomassie. To quantify vimentin cleavage, the relative intensities of the cleaved vimentin fragment bands were measured via densitometry, then normalized to the intensity of the full-length vimentin band present in the samples. To quantify changes in solubility, the relative intensities of the full-length vimentin bands in the detergent-soluble and insoluble fractions were measured via densitometry, then normalized to the intensity of the full-length vimentin band in the total cell lysates. The expression levels of calpain-1, calpain-2, and calpastatin were measured via densitometry. The graphical data were generated using the GraphPad Prism software and analyzed using multiple T-tests, one-way ANOVA, or 2-way ANOVA.

